# Can the location of a trophectoderm biopsy contribute to human blastocyst development ?

**DOI:** 10.1101/109298

**Authors:** Tomoe Takano, Miyako Funabiki, Sagiri Taguchi, Fumie Saji, Namiko Amano, Kate Young Louise, Yoshitaka Nakamura

**Affiliations:** Oak Clinic, Osaka and Tokyo, Japan 2-7-9 Tamade-Nishi, Nishinari-ku, Osaka, 557-0045

## Abstract

The influence of the location of a trophectoderm biopsy in human blastocysts on the development of those blastocysts has not yet been investigated. In our prospective study (n=92), our multivariate logistic regression analysis indicated that blastocoel development was influenced by the location of the trophectoderm biopsy (p=0.049) and by the type of human blastocyst used (fresh or thawed) (p=0.037), regardless of the patient’s age (p=0.507) and the number of days for the human blastocyst in the pre-trophectoderm biopsy (p=0.239). Therefore, when a trophectoderm biopsy is close to the inner cell mass (ICM) in human blastocysts, it improves the progress of blastocoel development.

Clinical evidence suggests that the progress of blastocoel development is a predictor of clinical outcomes after single blastocyst transfer. Therefore, when the trophectoderm biopsy is done from near the ICM, improvement of clinical outcomes after single blastocyst transfer may be expected.

## Introduction

Trophectoderm biopsy for human blastocysts is conducted for preimplantation genetic screening (PGS) and/or preimplantation genetic diagnosis (PGD) in many *in vitro* fertilization (IVF) clinics [1-3].

However, the influence of the location of a trophectoderm biopsy in human blastocysts on the development of those blastocysts has not yet been investigated.

## Materials and Methods

### Study design

The present study was an experimental study involving 92 patients (median age 34.3 years old) with infertility at our institute.

Discarded embryos used in the study were collected with the patients’ informed consent and were cultured. They were either fresh or frozen-thawed.

### The study protocol

Each patient was assigned to one of the following three treatment groups as follows (Fig. 1).

**Fig. 1:**
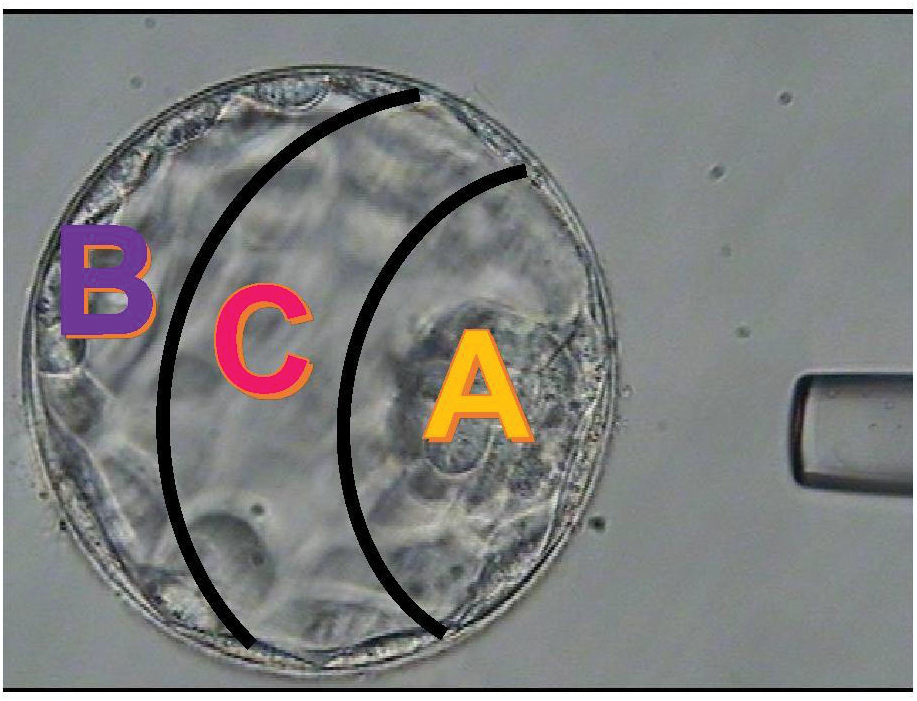
The location of a trophectoderm biopsy.

> Group A: location close to the inner cell mass (ICM) (n#x003D;29)
>
> Group B: location distant from the ICM (n=32)
>
> Group C: location between A and B (n=31)

The influence of the location of the trophectoderm biopsy within the human blastocysts on the development of those blastocysts was compared between pre- and post-trophectoderm biopsies, according to the Gardner and Schoolcraft scoring system ^4^. Human blastocyst development was scored by blastocoel stage (from 1 to 6: highest score is 6), ICM grade (highest score A, followed by B and C) and trophectoderm (TE) grade (highest score A, followed by B and C) [4]. Higher scores were considered improvements [4]. Two clinical embryologists evaluated the human blastocyst development.

The time between the post-trophectoderm biopsies and the evaluation of the development of the human blastocysts was 24 hours.

### Institutional Review Board (IRB) approval

This study was approved by the IRB of Oak Clinic, Osaka and Tokyo, Japan (the approval number is 2013081904).

The patients provided informed consent.

### Statistical analyses

Statistical tests were performed using Dr. SPSS II for Windows (SPSS Japan, Inc., Tokyo), and significance was defined as p<0.05. Statistical analyses of group differences were analyzed using Fisher’s exact test.

## Results

### The rate of blastocoels that showed developmental progress

According to the Gardner and Schoolcraft scoring system [4], degree (one up to six) of expansion of the blastocoel indicates progress of blastocoel development. The rate of blastocoels that showed developmental progress in Group A was significantly higher (p=0.024, Fisher’s exact test) than in Group B: 25/29 (86.2%) versus 19/32 (59.4%), respectively (Fig.2). The location of the trophectoderm biopsy in the human blastocysts did not change the trophectoderm and ICM grading.

**Fig. 2:**
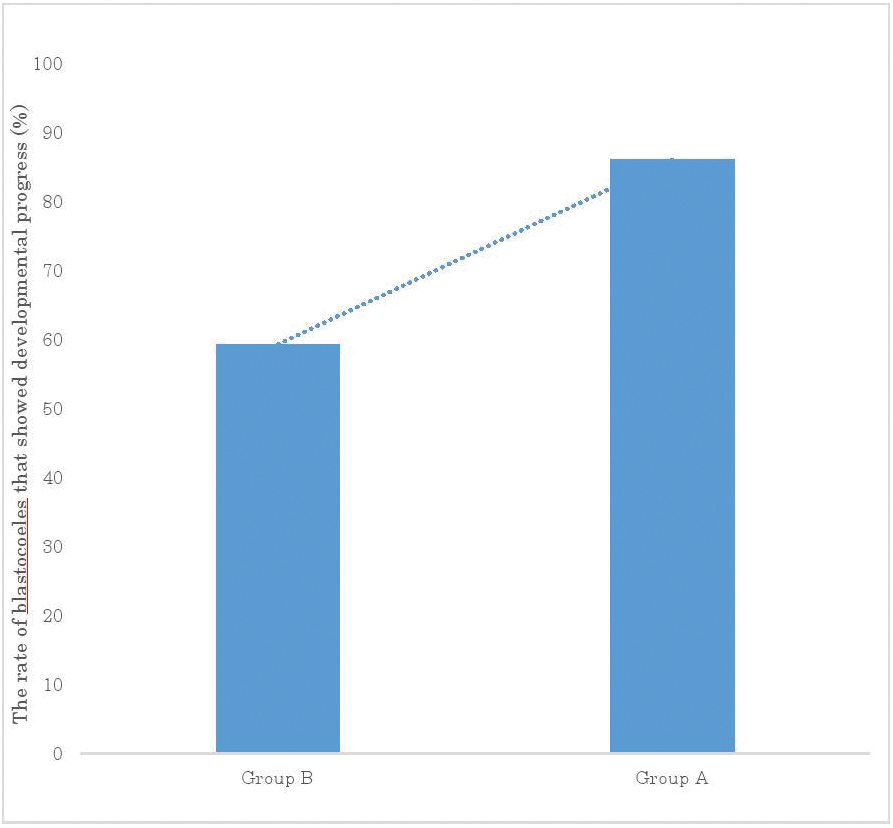
The rate of blastocoels that showed developmental progress.

### Multivariate logistic regression analysis

Multivariate logistic regression analysis (Table 1) indicated that blastocoel development was influenced by the location of the trophectoderm biopsy (p=0.049) and by the type of human blastocyst used (fresh or thawed) (p=0.037), regardless of the patient’s age (p=0.507) and the number of days for the human blastocyst in the pre-trophectoderm biopsy (p=0.239).

**Table 1:**
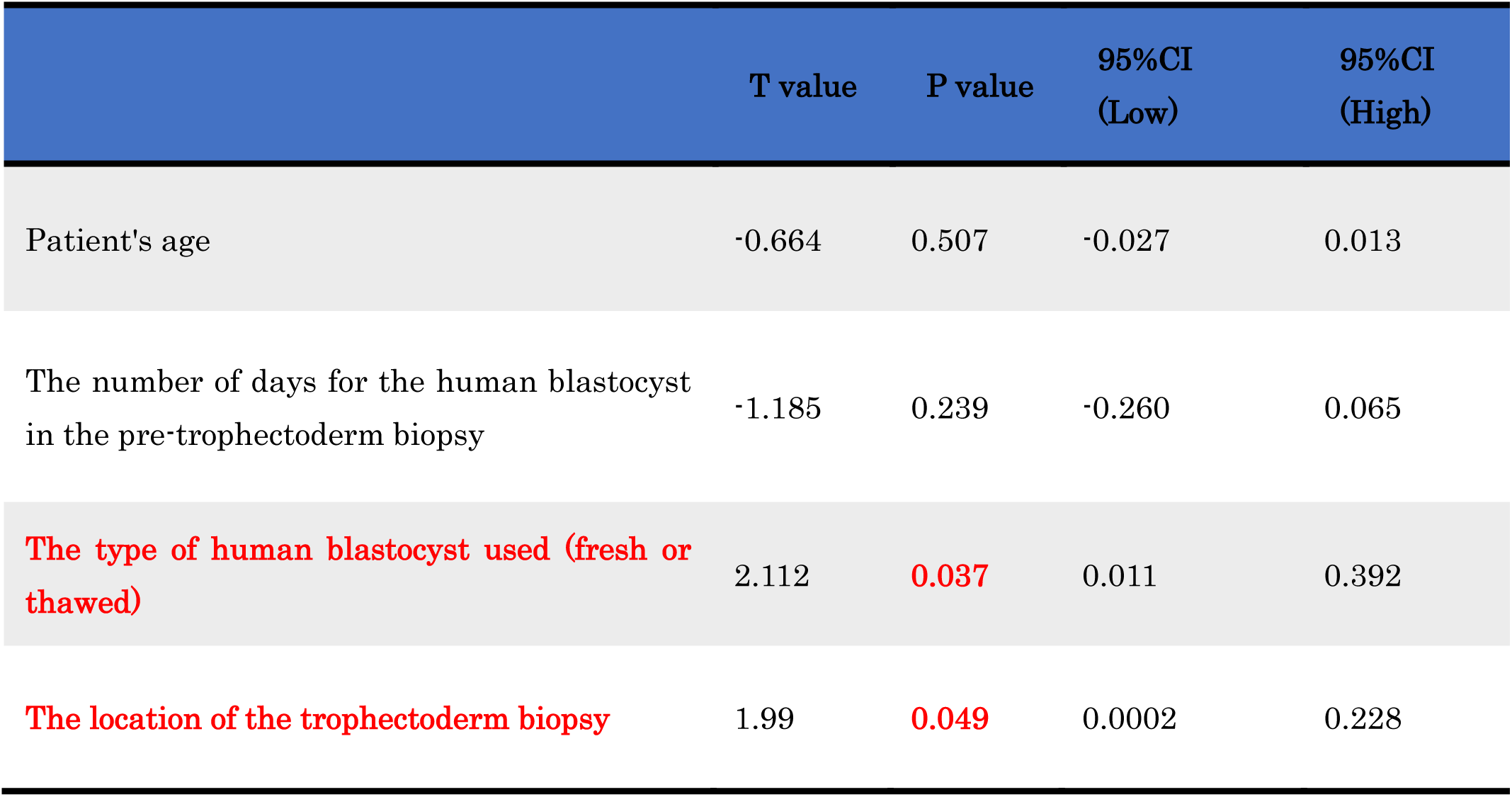
Multivariate logistic regression analysis.

|  | T value | P value | 95%CI<br>(Low) | 95%CI<br>(High) |
| --- | --- | --- | --- | --- |
| Patient's age | -0.664 | 0.507 | -0.027 | 0.013 |
| The number of days for the human blastocyst<br>in the pre-trophectoderm biopsy | -1.185 | 0.239 | -0.260 | 0.065 |
| **The type of human blastocyst used (fresh or**<br>**thawed)** | 2.112 | **0.037** | 0.011 | 0.392 |
| **The location of the trophectoderm biopsy** | 1.99 | **0.049** | 0.0002 | 0.228 |

95% CI means 95 % confidence interval.

## Discussion

The present study is the first to report that when a trophectoderm biopsy is close to the ICM in human blastocysts, it improves the progress of blastocoel development.

Clinical evidence suggests that the progress of blastocoel development is a predictor of clinical outcomes after single blastocyst transfer [5-8]. Therefore, when the trophectoderm biopsy is done from near the ICM, improvement of clinical outcomes after single blastocyst transfer may be expected.

However, when a trophectoderm biopsy is close to the ICM in human blastocysts, the risk is still unclear. Therefore, the risk and benefit in the clinical settings should be evaluated in the near future.

## Acknowledgements

We are grateful to the physicians, nurses and clinical embryologists for their assistance with the design of this study and/or the experiments performed at our clinic.

### Author contributions

T.T.: Conception and design of the study, provision of the study materials, collection and/or assembly of the data, analysis and interpretation of the data, writing of the manuscript, and final approval of the manuscript.

S.T.: Provision of the study materials, collection and/or assembly of the data, analysis and interpretation of the data and final approval of the manuscript.

M.F.: Provision of the study materials, collection and/or assembly of the data, analysis and interpretation of the data and final approval of the manuscript.

F.S.: Provision of the study materials, collection and/or assembly of the data and final approval of the manuscript.

N.A.: Provision of the study materials, collection and/or assembly of the data and final approval of the manuscript.

K. L.Y.: Provision of the study materials, collection and/or assembly of the data and final approval of the manuscript.

Y.N.: Analysis and interpretation of the data, writing of the manuscript, and final approval of the manuscript.

## References

1. Zakharova EE, Zaletova VV, Krivokharchenko, AS. Biopsy of human morula-stage embryos: outcome of 215 IVF/ICSI cycles with PGS. PLoS ONE. 2014; 9: e106433; 10.1371/journal.pone.0106433.

2. Gleicher N, Batad DH. A review of, and commentary on, the ongoing second clinical introduction of preimplantation genetic screening (PGS) to routine IVF practice. J Assist Reprod Genet. 2012; 29: 1159–1166.

3. Harper JC, Sengupta SB. Preimplantation genetic diagnosis: state of the art 2011. Hum Genet. 2012; 131: 175–186.

4. Gardner DK, Schoolcraft WB. *In vitro* culture of human blastocysts (eds. Jansen, R and Mortimer, D.), Parthenon Publishing London; 1999, pp. 378–388.

5. Du QY, Wang EY, Huang Y, Guo XY, Xiong YJ, Yu YP, et al. Blastocoele expansion degree predicts live birth after single blastocyst transfer for fresh and vitrified/warmed single blastocyst transfer cycles. Fertil Steril. 2016; 105: 910–919.

6. Thompson SM, Onwubalili N, Brown K, Jindal SK, McGovern PG. Blastocyst expansion score and trophectoderm morphology strongly predict successful clinical pregnancy and live birth following elective single embryo blastocyst transfer (eSET): a national study. J Assist Reprod Genet. 2013; 30: 1577–1581.

7. Van den Abbeel, E, Balaban B, Ziebe S, Lundin K, Cuesta MJ, Klein BM, et al. Association between blastocyst morphology and outcome of single blastocyst transfer. Reprod Biomed Online. 2013; 27: 353–361.

8. Ahlstrom A, Westin C, Wikland M, Hardarson T. Prediction of live birth in frozen-thawed single blastocyst transfer cycles by pre-freeze and postthaw morphology. Hum Reprod. 2013; 28: 1199–1209.

